## supporting information for "Predicting Relative Populations of Protein Conformations without a Physics Engine Using AlphaFold2"

### Supplementary Materials for “Predicting Relative Populations of Protein Conformations without a Physics Engine Using AlphaFold2"

**Gabriel Monteiro da Silva and Jennifer Y. Cui**
*Brown University Department of Molecular Biology, Cell Biology, and Biochemistry, Providence, RI, USA*

**David C. Dalgarno**
*Dalgarno Scientific LLC, Brookline, MA, USA*

**George P. Lisi and Brenda M. Rubenstein**
*Brown University Department of Molecular Biology, Cell Biology, and Biochemistry*
*Brown University Department of Chemistry*
*Providence, RI, USA*

July 27, 2023

### Abl1 Ortholog Sequences Used to Generate Multiple Sequence Alignments

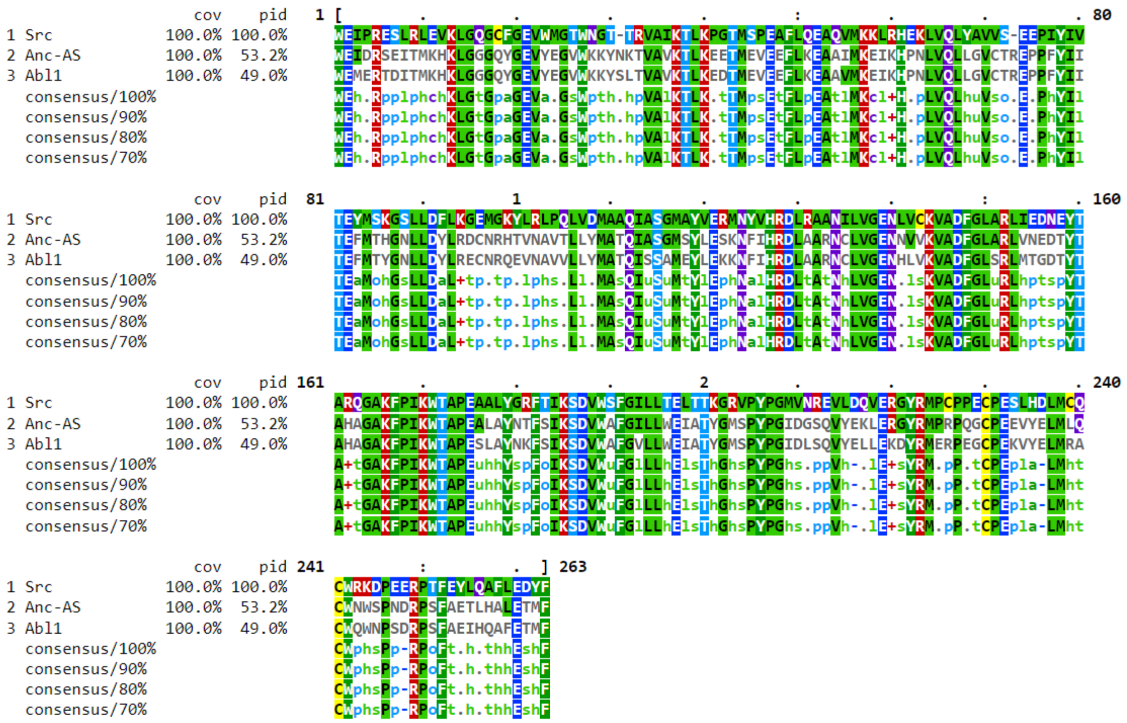

[width=450pt]

**Fig. S1:** Sequences of the Abl1, Src, and Anc-AS kinase cores used to generate MSAs and as input for subsampled AlphaFold 2.

### Comparisons Between AF2 Predictions and Molecular Dynamics Snapshots along the Ground to I2 Transition

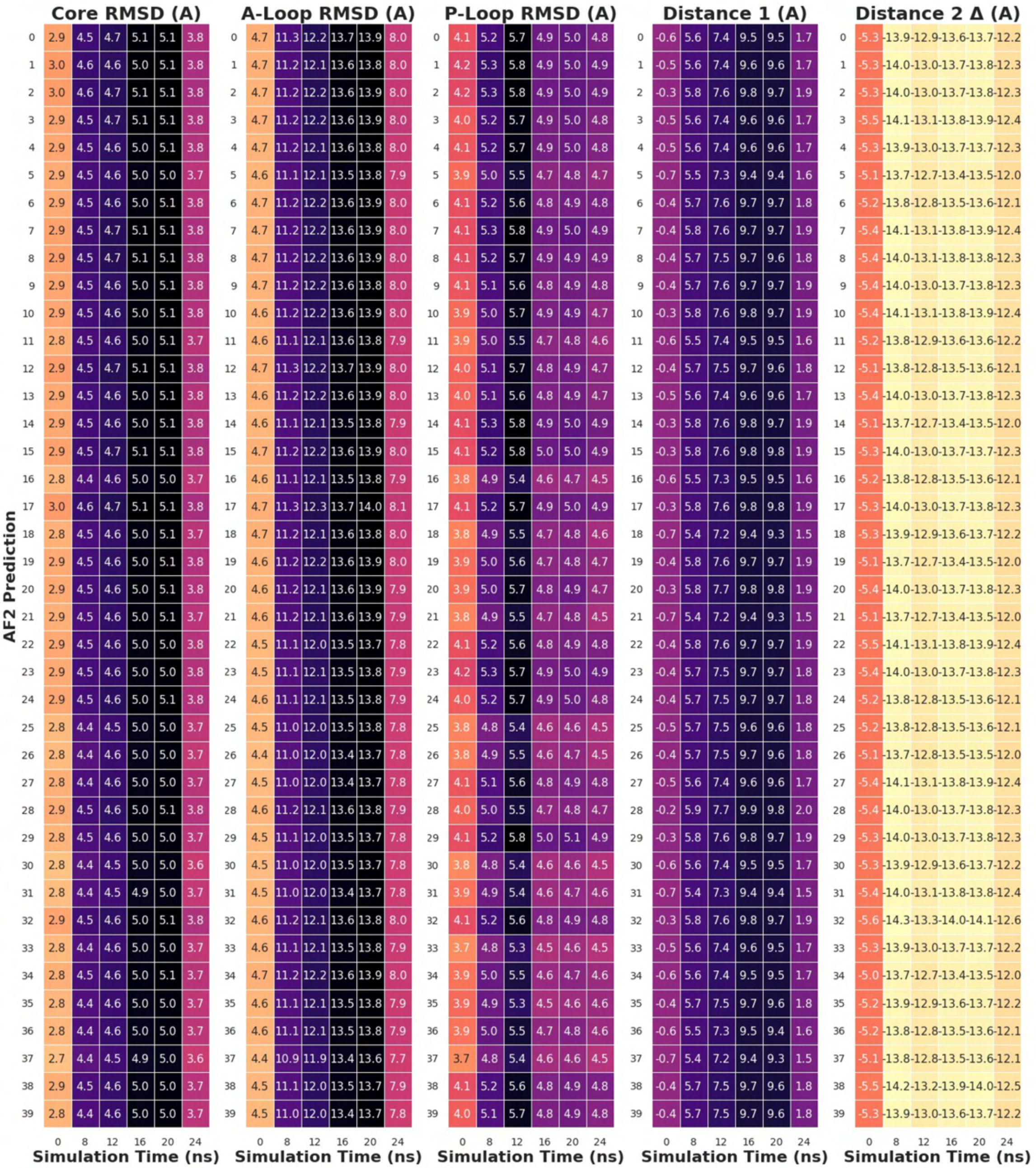

**Fig. S2:** Part one of four of the comparison between the values of five structural elements in the Abl1 kinase core known to change during the ground to I2 transition as measured from the ensemble of 160 subsampled AF2 predictions and six frames extracted from a molecular dynamics simulation trajectory spanning the transition at different time points. Core, P-Loop, and A-Loop RMSDs are defined as the backbone RMSDs of each AF2 prediction's kinase core (residues 242 to 459), activation loop (residues 379 to 395), or phosphate-binding loop (residues 244 to 256) vs. the kinase core, phosphate-binding loop, or activation loop backbone of the MD snapshot selected at each time point. Distance deltas are defined as the difference in atom pair distances between each AF2 prediction and its respective MD snapshot. Distance 1 corresponds to the distance between the backbone oxygens of E377 and L409, and Distance 2 corresponds to the distance between the backbone oxygens of L409 and G457.

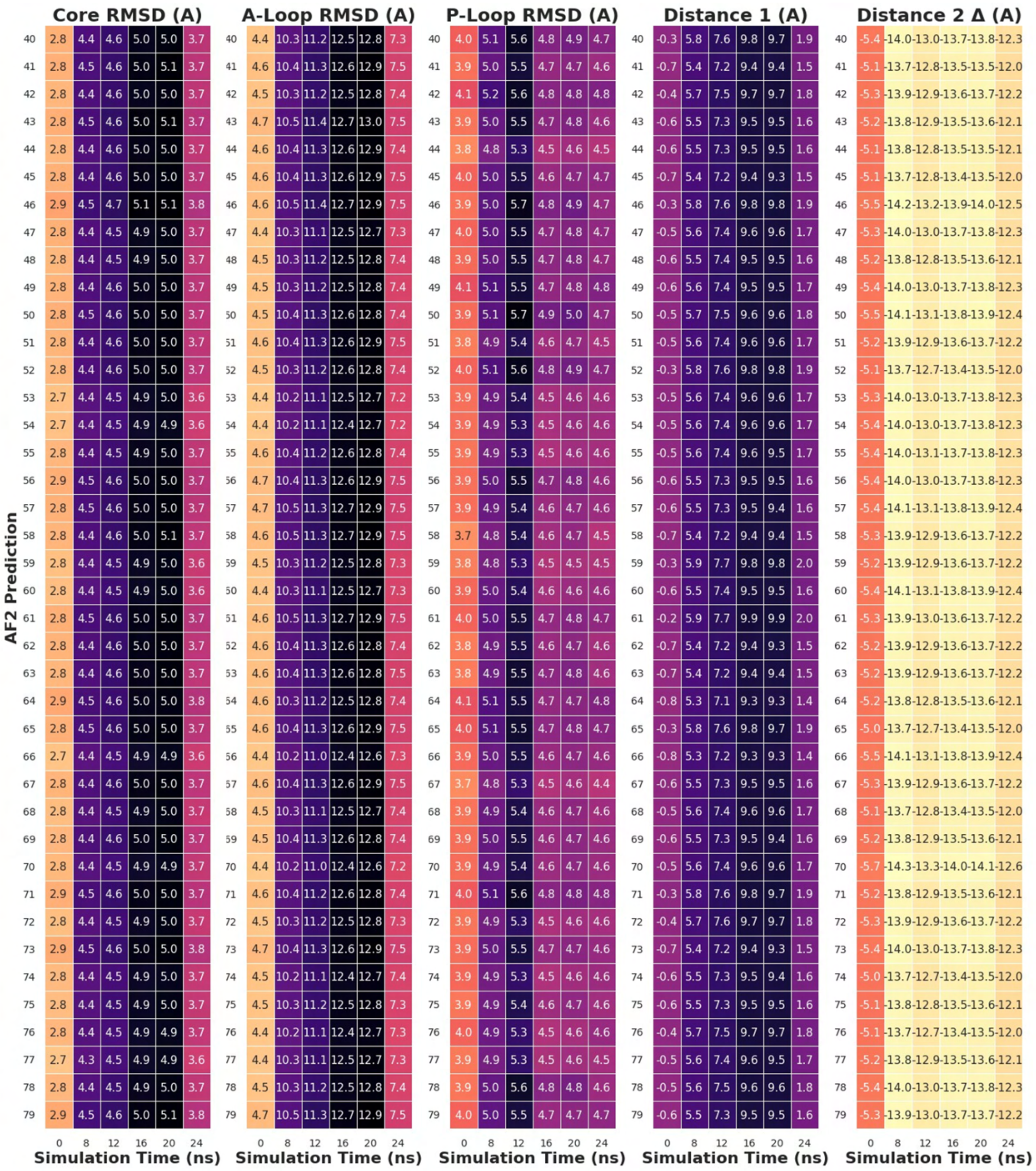

**Fig. S3:** Part two of four of the comparison between the values of five structural elements in the Abl1 kinase core known to change during the ground to I2 transition as measured from the ensemble of 160 subsampled AF2 predictions and six frames extracted from a molecular dynamics simulation trajectory spanning the transition at different time points. Core, P-Loop, and A-Loop RMSDs are defined as the backbone RMSDs of each AF2 prediction's kinase core (residues 242 to 459), activation loop (residues 379 to 395), or phosphate-binding loop (residues 244 to 256) vs. the kinase core, phosphate-binding loop, or activation loop backbone of the MD snapshot selected at each time point. Distance deltas are defined as the difference in atom pair distances between each AF2 prediction and its respective MD snapshot. Distance 1 corresponds to the distance between the backbone oxygens of E377 and L409, and Distance 2 corresponds to the distance between the backbone oxygens of L409 and G457.

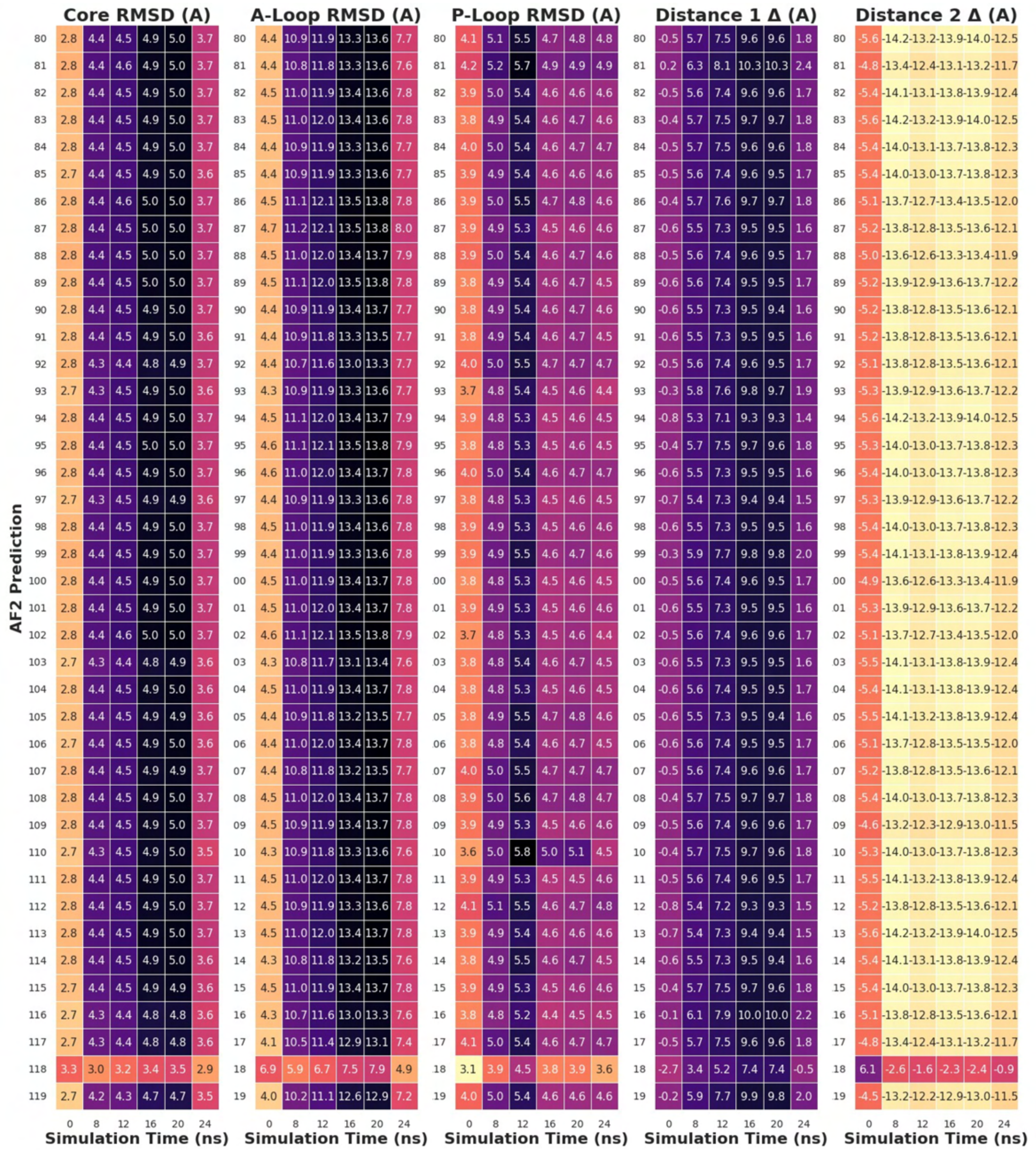

**Fig. S4:** Part three of four of the comparison between the values of five structural elements in the Abl1 kinase core known to change during the ground to I2 transition as measured from the ensemble of 160 subsampled AF2 predictions and six frames extracted from a molecular dynamics simulation trajectory spanning the transition at different time points. Core, P-Loop, and A-Loop RMSDs are defined as the backbone RMSDs of each AF2 prediction's kinase core (residues 242 to 459), activation loop (residues 379 to 395), or phosphate-binding loop (residues 244 to 256) vs. the kinase core, phosphate-binding loop, or activation loop backbone of the MD snapshot selected at each time point. Distance deltas are defined as the difference in atom pair distances between each AF2 prediction and its respective MD snapshot. Distance 1 corresponds to the distance between the backbone oxygens of E377 and L409, and Distance 2 corresponds to the distance between the backbone oxygens of L409 and G457.

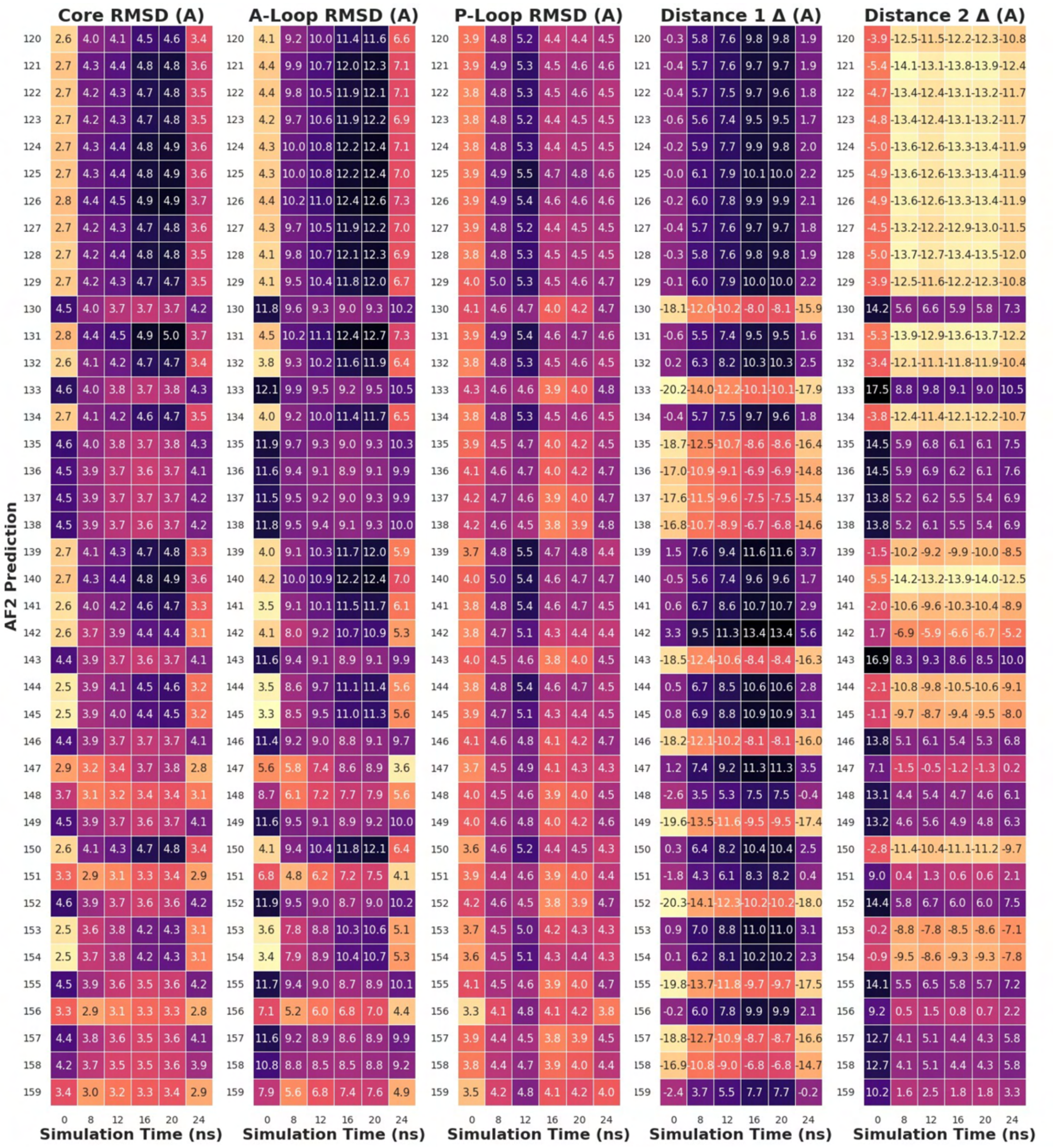

**Fig. S5:** Part four of four of the comparison between the values of five structural elements in the Abl1 kinase core known to change during the ground to I2 transition as measured from the ensemble of 160 subsampled AF2 predictions and six frames extracted from a molecular dynamics simulation trajectory spanning the transition at different time points. Core, P-Loop, and A-Loop RMSDs are defined as the backbone RMSDs of each AF2 prediction's kinase core (residues 242 to 459), activation loop (residues 379 to 395), or phosphate-binding loop (residues 244 to 256) vs. the kinase core, phosphate-binding loop, or activation loop backbone of the MD snapshot selected at each time point. Distance deltas are defined as the difference in atom pair distances between each AF2 prediction and its respective MD snapshot. Distance 1 corresponds to the distance between the backbone oxygens of E377 and L409, and Distance 2 corresponds to the distance between the backbone oxygens of L409 and G457.

### GMCSF Chemical Shift Perturbations

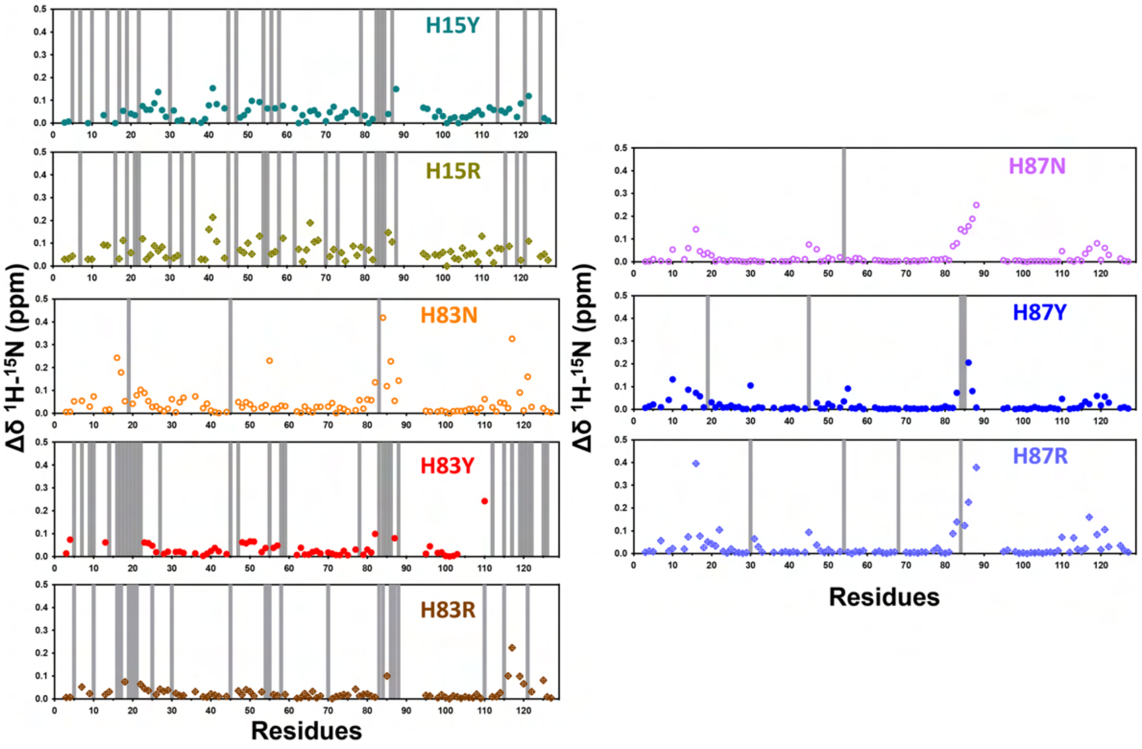

**Fig. S6:** N15-H1 Chemical shift perturbations for mutant GMCSF constructs in reference to wild-type GMCSF peaks. Vertical bars indicate residues where the signal was lost due to chemical exchange broadening.

### Optimization of AF2 Parameters for the GMCSF Protein

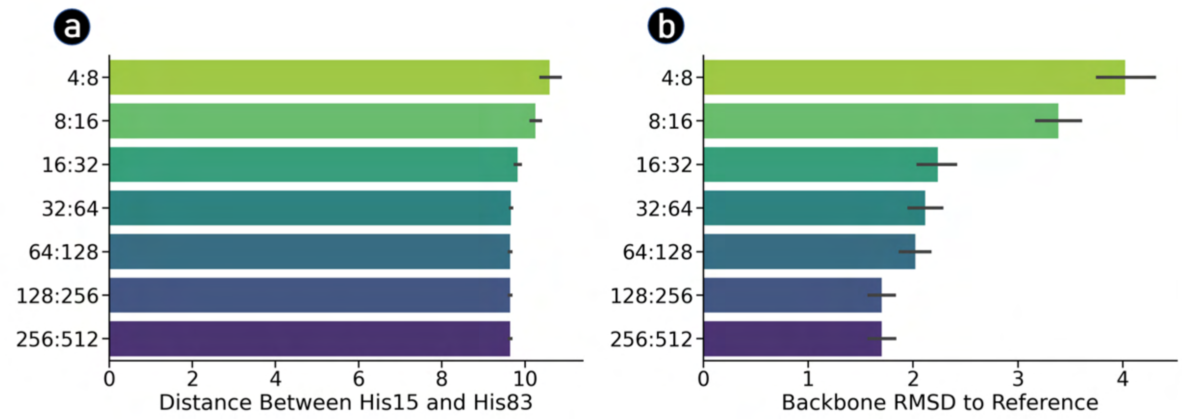

**Fig. S7:** Optimal AF2 subsampling parameters for GMCSF. (a) Effects of modifying the *max_seqs* and *extra_seqs* values on the diversity of the distances between the H15 and H83 residues observed, which is a proxy for the opening of the heparin-binding site in GMCSF. (b) Effects of modifying the *max_seqs* and *extra_seqs* values on the diversity of the root mean square deviation of atomic positions (RMSD) of the GMCSF backbone with respect to the ground state reference (PDB 1CSG).

### GMCSF Dynamics Prediction

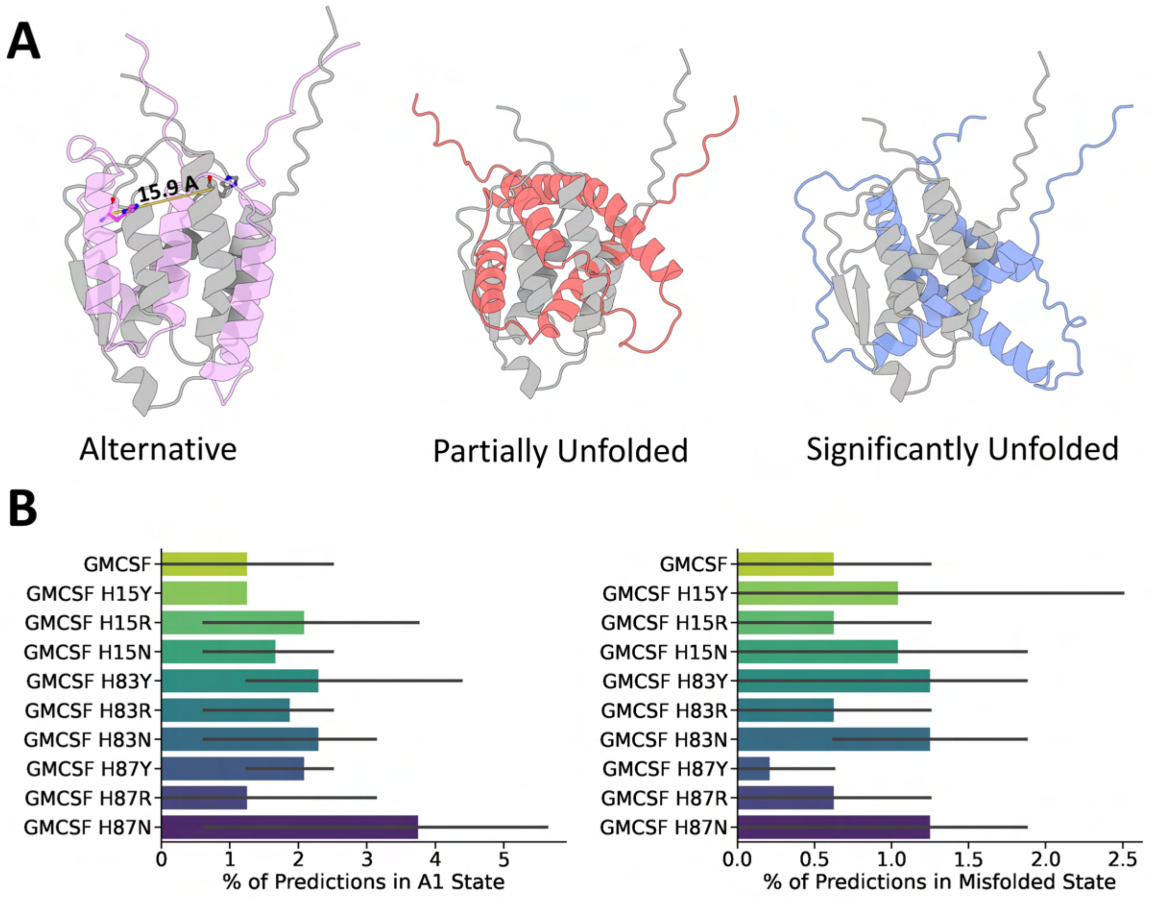

**Fig. S8:** Unusual GMCSF states predicted by subsampled AF2 and the respective populations of those states. (a) Structure of the most common alternative state predicted by AF2 (A1, in pink) aligned with and overlain on a ground state prediction (in grey). The distance between H83 in the reference and in conformation A1 is displayed as a measure of the difference between the conformations. Also shown are two misfolded/unfolded predictions aligned with and overlain on the ground state prediction (in grey). (b) AF2 predictions of the relative populations of the A1 conformation and the misfolded/unfolded structures. Conformations were classified as the A1 conformation based on the distance between the H15 and H83 residues (greater than 11 Å) and overall backbone RMSD vs. ground state reference (greater than 5 A but less than 10 Å), while they were classifies as misfolded/unfolded based on overall backbone RMSD vs. the ground state reference (equal to or greater than 13 Å).

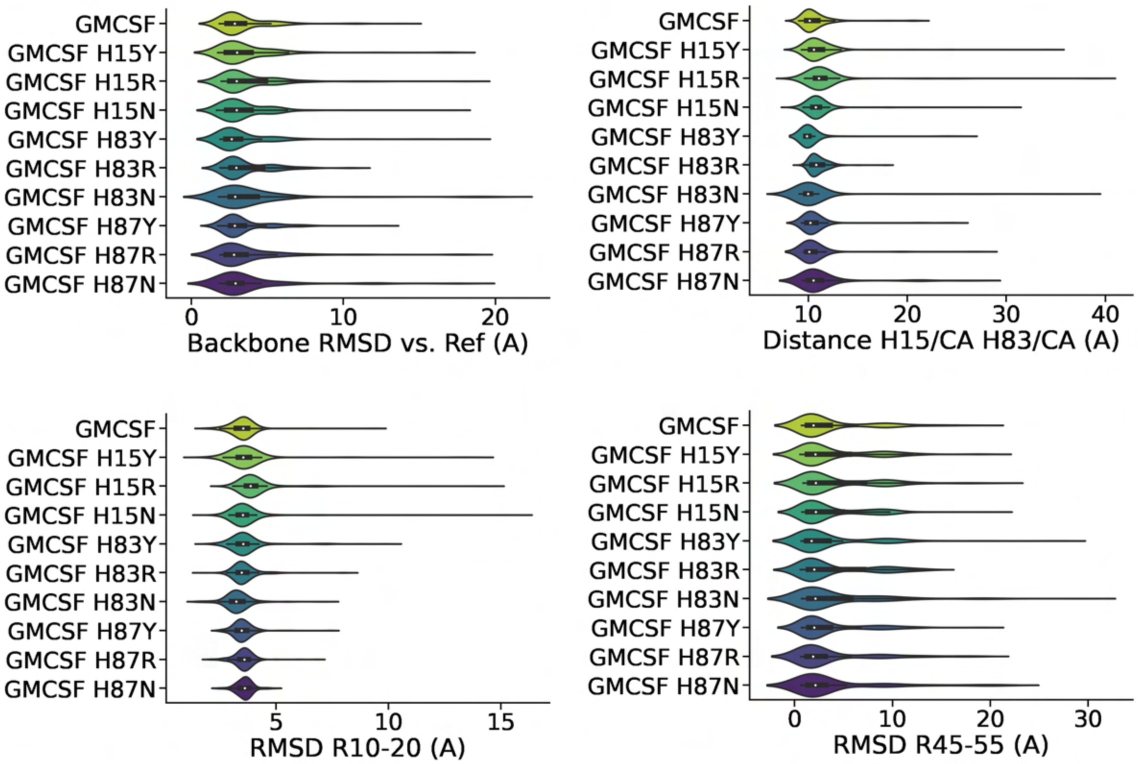

**Fig. S9:** AF2 predictions of the distributions of different GMCSF properties. Every RMSD measurement was taken with respect to the ground state reference (PDB 1CSG).

### Optimization of AF2 Parameters for the Abl1 Protein

**Table S1:** Optimized AF2 parameters for predicting Abl1 ensembles.

| parameter_test | max_seq | extra_seq | n_recycles | n_models | n_seeds | %_notground |
|---|---|---|---|---|---|---|
| **t_max_extra_1** | 32 | 64 | 4 | 5 | 32 | 2 |
| **t_max_extra_2** | 64 | 128 | 4 | 5 | 32 | 5 |
| **t_max_extra_3** | 128 | 256 | 4 | 5 | 32 | 9 |
| **t_max_extra_4** | 256 | 512 | 4 | 5 | 32 | 18 |
| **t_max_extra_5** | 512 | 1024 | 4 | 5 | 32 | 15 |
| **t_max_extra_6** | 2048 | 4096 | 4 | 5 | 32 | 7 |
| **t_max_extra_7** | 4098 | 8192 | 4 | 5 | 32 | 6 |
| **t_max_extra_8** | 512 | 32 | 4 | 5 | 32 | 18 |
| **t_max_extra_9** | 32 | 512 | 4 | 5 | 32 | 1 |
| **t_nseeds_1** | 256 | 512 | 4 | 5 | 128 | 12 |
| **t_nseeds_2** | 256 | 512 | 4 | 5 | 300 | 12 |
| **t_nrecycles_1** | 32 | 64 | 8 | 5 | 128 | 0 |
| **t_nrecycles_2** | 32 | 64 | 8 (kept) | 5 | 128 | 2 |
| **t_nrecycles_3** | 256 | 512 | 8 | 5 | 128 | 8 |
| **t_nrecycles_4** | 256 | 512 | 8 (kept) | 5 | 128 | 21 |

### AF2 Predictions of the Relative State Populations of Abl1 Kinase Core Mutants

**Table S2:** Abl1 kinase core mutants and their observed or expected effects on the relative populations of the active (Ground), inactive 1 (I1), or inactive 2 (I2) states.

| | Ground | I1 | I2 |
|---|---|---|---|
| **Wild-Type** | 88 | 6 | 6 |
| **M290L** | 55 | 10 | 35 |
| **L301I** | 25 | 10 | 65 |
| **M290L + L301I** | 8 | 10 | 82 |
| **F382L** | 90 | 0 | 10 |
| **F382Y** | 10 | 0 | 90 |
| **F382V** | 5 | 0 | 95 |
| **I2M** | 10 | 0 | 90 |
| **E255V (I2M background)** | nr | nr | 45 |
| **T315I (I2M background)** | 93 | 0 | 7 |
| **E255V + T315i** | nr | nr | nr |

### Molecular Dynamics and WESTPA2 Simulations

Molecular dynamics simulations of wild-type Abl1 were conducted using the OpenMM software package [1] with the amber99sb-ildn force field [2] and the tip3p water model [3] at 300 K and 1 atm. The lowest energy Abl1 structure from the PDB 6XR6 NMR ensemble was solvated within a dodecahedron box and charges were neutralized by replacing a number of solvent atoms with chloride and sodium ions. Following solvation, we minimized the energy of each system using a steepest-descent algorithm until the maximum force on any given atom was less than 1000 kJ/mol/min or until 50,000 minimization steps were conducted. We ran the simulations with a 1 fs time step during the equilibration phase and a 2 fs time step during the production phase. We equilibrated solvent atoms first for 1 ns in the NVT ensemble and then for 1 ns in the NPT ensemble with solute heavy atoms restrained using the LINCS algorithm with a spring constant of 1,000 kJ/mol/$m^2$ [4]. The production phase (in the NPT ensemble) followed the equilibration phase but without restraints.

We used the WESTPA2 [5] enhanced-sampling method to access the timescales necessary to simulate the inactivation pathway of Abl1. This was done via two WESTPA2 simulations (ground to I1 and I1 to I2). As progress coordinates for the ground to I1 transition, we defined the distance between the backbone oxygen of V299 and the center of mass of the carboxyl group of D381 as PC1; and the angle formed by the center of mass of the carboxyl group of D381, the backbone oxygen of K379, and the center of mass of the aromatic ring of F382 as PC2. For the I1 to I2 transition, we defined the distance between the backbone oxygen of L409 and the backbone oxygen of E377 as PC1; and the distance between backbone oxygen of L409 and the backbone oxygen of G4598 as PC2. Representative illustrations of the progress coordinates used in this protocol are in Figure S10, and their distributions and start/end state definitions are described in Fig S11. We ran WESTPA2 for 300 iterations for each leg of the transition, with the number of walkers per iteration varying from 64 to 512 due to the adaptive binning scheme, and 100 ps per iteration, totaling over 9 us of aggregate simulation time for each leg of the transition.

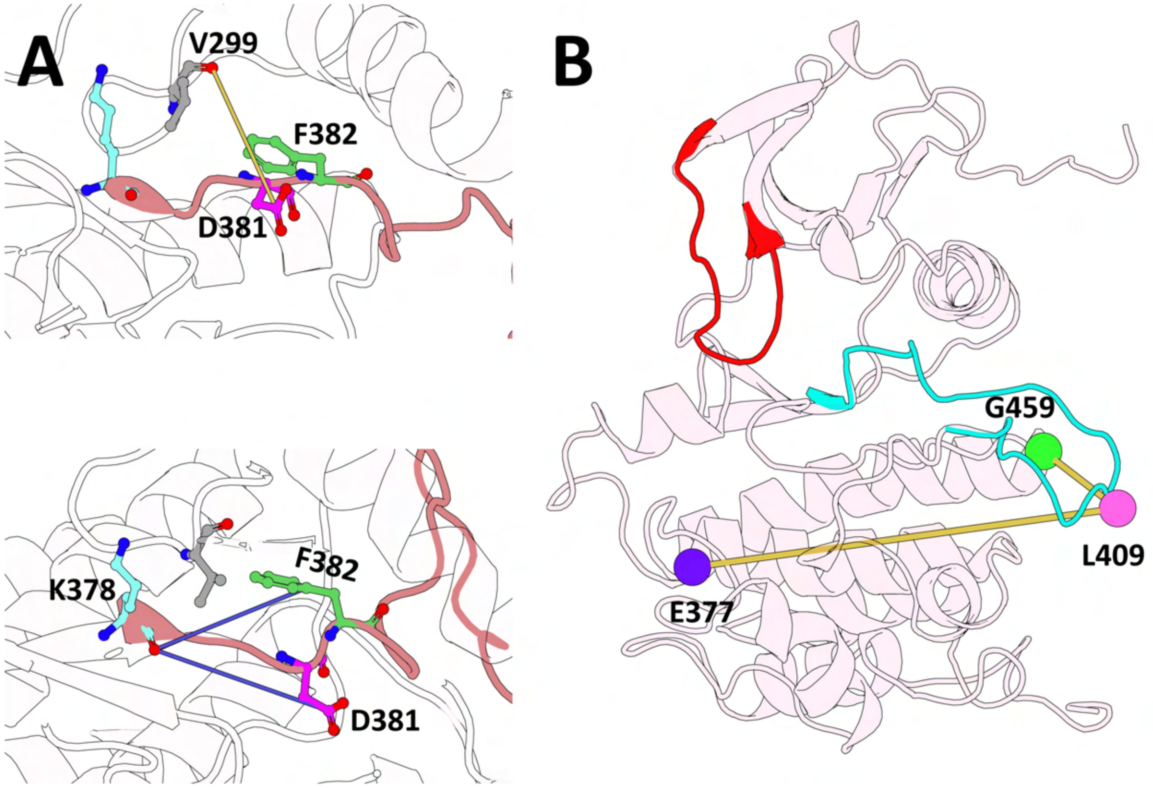

**Fig. S10:** Progress coordinates used in the WESTPA2 simulations of wild-type Abl1. (A) Progress coordinates used in sampling the transition from the ground to the I1 state. (B) Progress coordinates used in sampling the transition from the ground to the I2 state.

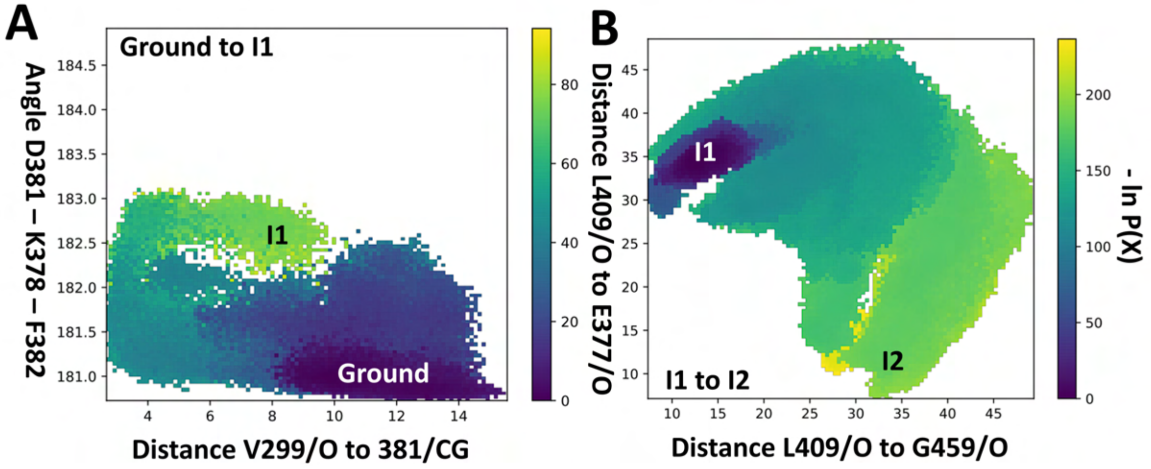

**Fig. S11:** Distribution of values for the progress coordinates used in either the transition from the (A) ground to the I1 state or (B) I1 to I2 state.

### References

1. Eastman, P. *et al.* OpenMM 7: Rapid development of high performance algorithms for molecular dynamics. *PLOS Computational Biology* **13** (ed Gentleman, R.) e1005659 (July 2017).
2. Lindorff-Larsen, K. *et al.* Improved side-chain torsion potentials for the Amber ff99SB protein force field. *Proteins: Structure, Function, and Bioinformatics* **78,** 1950–1958 (Mar. 2010).
3. Mark, P. & Nilsson, L. Structure and Dynamics of the TIP3P, SPC, and SPC/E Water Models at 298 K. *The Journal of Physical Chemistry A* **105,** 9954–9960 (Oct. 2001).
4. Hess, B., Bekker, H., Berendsen, H. J. C. & Fraaije, J. G. E. M. LINCS: A linear constraint solver for molecular simulations. *Journal of Computational Chemistry* **18,** 1463–1472 (Sept. 1997).
5. Bogetti, A. T. *et al.* A Suite of Advanced Tutorials for the WESTPA 2.0 Rare-Events Sampling Software [Article v2.0]. *Living Journal of Computational Molecular Science* **5** (2022).
